# Experimental and Simulated Approaches to Roadkill Persistence: Implications for Road Mortality Assessment

**DOI:** 10.1101/2025.07.15.664867

**Authors:** Annaëlle Bénard, Thierry Lengagne

## Abstract

The time during which roadkill remains available for detection following a collision is a central issue in road ecology. Not accounting for roadkill persistence leads to underestimating the actual impact of vehicular traffic on populations. Moreover, disregarding the inconsistent rates of disappearance across species, seasons and roads prevents carcass counts from being comparable both within and across studies. This can lead to a misidentification of roadkill hotspots and species for which road mortality is the highest. However, finding the appropriate experimental method to estimate roadkill persistence is difficult: both placing carcasses on roads to conduct removal trials and periodic monitoring of roadkill already present may fail to accurately describe the underlying distribution in persistence times. In this study, in order to assess the potential for systematic bias in roadkill persistence studies depending on the methodology, we simulate roadkill appearance and removal from the road by fitting a survival distribution based on experimental data. We then explore the discrepancies between methods by analysing empirical persistence data derived from both carcass trials and periodic monitoring. We find that, both in theory and in practice, monitoring tends to produce inflated estimates of roadkill persistence compared to carcass removal trials. Accurately describing the distribution of roadkill persistence times may therefore present a greater challenge than previously anticipated.

## INTRODUCTION

The ever-expanding global road network encompasses all the major drivers of biodiversity loss identified by the IPBES (Díaz et al., 2019). Terrestrial transportation contributes to global greenhouse gases emissions associated to climate change (European Environment Agency, 2024), while roadsides suffer from noise pollution (Parris & Schneider, 2009), light pollution (Gaston & Holt, 2018) and accumulate chemical pollutants such as heavy metals and salts (Hwang et al., 2016), impacting the fauna and flora of the surrounding habitats (Seiler, 2001). Road infrastructures and road traffic contribute to habitat fragmentation (Holderegger & Di Giulio, 2010). By disrupting ecological processes, terrestrial transportation is therefore detrimental to the survival of populations for many species (Forman & Alexander, 1998). At last, collisions between wildlife and terrestrial transports may be a substantial source of mortality (Moore et al., 2023) amounting to an estimated 220 million birds and mammals killed in Europe annually (Grilo et al., 2020).

Road mortality is often studied by performing standardized surveys of the carcasses present on roads. The data often serves as a basis to identify collisions hot-spots (Girardet et al., 2015; Gomes et al., 2009; Teixeira, Coelho, Esperandio, Rosa Oliveira, et al., 2013) and implement mitigation measures such as crossing structures and fences (Aresco, 2003; Rytwinski et al., 2016). The raw number of detected carcasses is, however, an under-estimation of the real number of wildlife-vehicle collisions as a result of imperfect carcass detection during surveys and limited carcass persistence on roads (Teixeira, Coelho, Esperandio, & Kindel, 2013). Carcass detectability is strongly influenced by the size of the animal (Barrientos et al., 2018), the position of the carcass on the road (R. A. L. Santos & Ascensão, 2019) and the travel speed and level of experience of the observers (Collinson et al., 2014; Guinard et al., 2023; Langen et al., 2007; Paraguassu-Chaves et al., 2020). In addition to imperfect detection, the persistence of carcasses on roads following a collision presents an additional challenge: roadkill surveys that are insufficiently frequent compared to the persistence of roadkill will result in an underestimation of the occurrence of wildlife-vehicle collisions (Henry et al., 2021) and can lead to a misidentification of roadkill hotspots (S. M. Santos et al., 2015).

As roadkill usually persists for a few days (Franceschi et al., 2021; Gerow et al., 2010; S. M. Santos et al., 2011; Winton et al., 2018) or hours (Beckmann & Shine, 2015; Bénard et al., 2024; Cabrera-Casas et al., 2020; Stewart, 1971), a significant proportion of studies—which rely on roadkill data obtained through bi-weekly, weekly or monthly surveys—would indeed miss an important part of the road mortality. In addition to underestimating the mortality, differences in searcher efficiency, carcass persistence and survey frequencies makes it impossible to compare raw mortality counts across studies (Huso, 2011). Studies show that roadkill persistence is a complex issue, influenced by traffic, season, weather, and species (Antworth et al., 2005; Ratton et al., 2014; R. A. L. Santos et al., 2016; S. M. Santos et al., 2011; Slater, 2002). Therefore, misidentification of certain seasons, road segments, or species as having higher collision rates may occur simply because they have longer carcass persistence (Menger et al., 2023), potentially resulting in misguided mitigation measures (Franceschi et al., 2021).

The issue of carcass persistence bias extends beyond roadkill monitoring and was initially examined in the context of mortality surveys in open fields, such as wind farms (Péron et al., 2013; Shoenfeld, 2004). The proposed corrective models for carcass counts typically necessitate unbiased empirical estimates of carcass persistence (Etterson, 2013; Huso, 2011; Teixeira, Coelho, Esperandio, & Kindel, 2013). For roads, several authors have included measures of persistence alongside the roadkill counting process to enable immediate adjustments to mortality rates (Boves & Belthoff, 2012; Gerow et al., 2010; Guinard et al., 2012; Teixeira, Coelho, Esperandio, & Kindel, 2013). Carcass persistence studies typically use one of two experimental approaches: (1) assessing the persistence of carcasses intentionally positioned on roadways, which serve as surrogates for actual fatalities (sometimes referred to as *carcass trials*), or (2) conducting regular surveys to identify new carcasses and subsequently document their presence during each monitoring event.

The first approach (carcass trials) may introduce several recognized biases, particularly if surrogate carcasses fail to accurately represent roadkill. For example, freezing carcasses during storage (Boves & Belthoff, 2012; Cabrera-Casas et al., 2020) may affect the scavenging rates compared to fresh roadkill (Etterson, 2013, but see (Barrientos et al., 2018). The location of the carcasses may also be significant, as scavengers expect to find carrion in specific areas and will remove carcasses located elsewhere at different rates (Bracey et al., 2016). Furthermore, analyses should account for the position of the carcass placed on the roadway—whether it is in the lane, in the path of vehicle tires, or on the verge—as it can impact its persistence (Bénard et al., 2024; R. A. L. Santos & Ascensão, 2019). Finally, the timing of placement (which simulates the time of collision) may also influence the resulting persistence estimates (Hubbard & Chalfoun, 2012).

However, the second experimental approach, as used for example by (Gerow et al., 2010; Henry et al., 2021; Ogletree & Mead, 2020; S. M. Santos et al., 2011), may also present inherent biases. Roadkill detection via surveys may over-represent larger individuals (Barrientos et al., 2018), or animal that are generally more easily detected due to their colour or placement on the road, resulting in biased samples that do not accurately represent the population of carcasses. Additionally, depending on the distribution of roadkill persistence times, the samples may suffer from length-bias, *i.e.,* that the probability of entering the study may be proportional to the length of the persistence itself (Shen et al., 2017). For example, if a significant proportion of carcasses persist for short periods of time (compared to the frequency of periodic monitoring), they will be under-represented in the sampling process because their presence on the road will often fall entirely between monitoring events. Conversely, carcasses may persist for extended periods (*e.g.,* located on road verges, where they can’t be run over by vehicles, and disfigured by traffic or hardened to the point of being unappealing to scavengers, (S. M. Santos et al., 2011). These animals will represent the tail of the persistence times distribution, which may be over-sampled during surveys, thereby skewing general roadkill persistence estimates upwards (Nielsen & Lang, 1999).

In the literature, studies using road surveys to recruit new roadkill carcasses into persistence experiments typically conduct these surveys between once a day (*e.g.,* S. M. Santos et al., 2011) and once a week (*e.g.,* Taylor & Goldingay, 2004). In contrast, studies that continuously or frequently monitor the disappearance of surrogate roadkill show that a significant portion of roadkill can disappear within 5 hours or less, sometimes even in a few minutes for small-bodied species (Beckmann & Shine, 2015; Bénard et al., 2024). Table 1 summarizes the current roadkill persistence estimates and experimental approaches to the best of our knowledge. This discrepancy between the frequency of surveys for new roadkill and the persistence of some animals suggests that, in some cases, estimates of roadkill persistence in studies recruiting carcasses directly from the road may be positively biased compared to studies placing and monitoring surrogate carcasses on roadways.

**Table 1:** Available estimates of the persistence of roadkill. Taxa: species in the sample; Method: the experimental approach, either monitoring roadkill discovered during periodic road surveys (D), or intentionally placing carcasses to simulate roadkill (P); Carcass search frequency: for studies using the “D” method, the frequency of road surveys to recruit new roadkill into the study; Model: statistical method, if any, to estimate roadkill persistence (Time-to-event: survival analysis, such as Cox-PH or Kaplan-Meier methods, CMR: Capture-Mark-Recapture); Measured Persistence: if available, the median or mean persistence, or if unreported, the proportion of carcasses remaining on roads after a set time interval.

| Study | Taxa | Method | Carcass search frequency | Model | Measured Persistence |
| --- | --- | --- | --- | --- | --- |
| Stewart, 1971 | House sparrow | P | - | - | 0% after 2 (highway) to 24 hours (country road) |
| Slater, 2002 | Domestic chicken | P | - | - | 1 – 5 hours (median<sup>3</sup>) |
| Taylor & Goldingay, 2004 | Mammals, Birds | D | Once a week | - | 39% after 7 days |
| Antworth et al., 2005 | Domestic chicken, Reptiles | P | - | - | 4 - 17 hours (median<sup>3</sup>) |
| Gerow et al., 2010 | All<sup>1</sup> | D | Once a day | MLE<sup>2</sup> | 3 days (mean) |
| Boves & Belthoff, 2012 | Barn owl | P | - | - | 100% after 3 days |
| Brzeziński et al., 2012 | Amphibians | P | - | Time-to-event | 41% after 24 hours |
| Guinard et al., 2012 | Birds | D and P | Every 6 to 18h | CMR | 54 - 67% after 10 days |
| Hubbard & Chalfoun, 2012 | Rattlesnake | P | - | - | 22 – 72% after 60 hours |
| Teixeira et al., 2013 | All<sup>1</sup> | D | Unknown (10 surveys/year) | - | 35 - 79% after 24h |
| Ratton et al., 2014 | Domestic chicken | P | - | - | 11% after 24 hours |
| Beckmann & Shine, 2015 | Amphibians | P | - | - | 23 minutes – 9 hours (mean) |
| Ruiz-Capillas et al., 2015 | Mammals | P | - | Fitting neg. exp. to the data | 7 days (median<sup>3</sup>) |
| R. A. L. Santos et al., 2016 | All<sup>1</sup> | D | Once a day | Time-to-event | 1 day (median<sup>3</sup>) |
| Winton et al., 2018 | Rattlesnake | P | - | - | 2 days (median<sup>3</sup>) |
| R. A. L. Santos & Ascensão, 2019 | Mammals | P | - | Time-to-event | 1 day (median<sup>3</sup>) |
| Ogletree & Mead, 2020 | All<sup>1</sup> | D | Twice a week | - | 1 – 3 days (mean) |
| Cabrera-Casas et al., 2020 | Reptiles | P | - | Time-to-event | 7 hours (median<sup>3</sup>) |
| Henry et al., 2021 | All<sup>1</sup> | D | Once a day | Time-to-event | 2 days (median<sup>3</sup>) |
| Franceschi et al., 2021 | Mammals | P | - | Time-to-event | 3 days (median<sup>3</sup>) |
| Bénard et al., 2024 | Amphibians, Birds | P | - | Time-to-event | 22 minutes – 18 hours (median<sup>3</sup>) |

*1. Mammals, birds, reptiles, amphibians. 2. Maximum Likelihood Estimator accounting for right-censored data. 3. The median persistence is defined as the time needed for persistence probabilities to drop below 50%.*

**Table 2:**
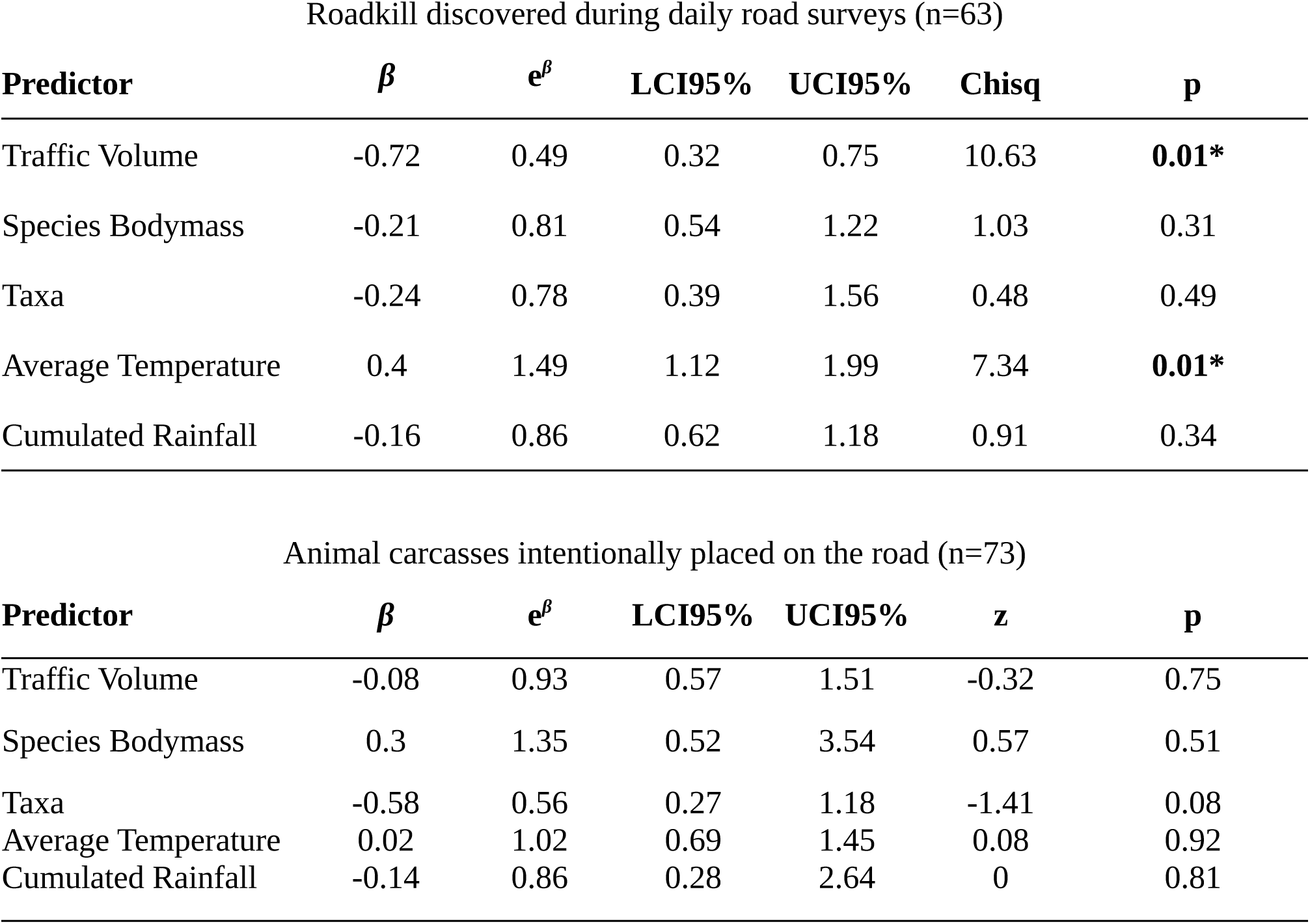
Cox-PH survival models of animal persistence on the road depending on the experimental approach: discovering and recording the presence of actual roadkill with daily road monitoring, or intentionally placing fresh animal carcasses on roads equipped with camera traps to monitor the survival continuously.

To assess the potential for length-biased sampling in roadkill persistence monitoring, we placed freshly deceased carcasses of various mammalian and avian species, representative of the local diversity, on roads and monitored their removal using camera traps. This method allows for precise and continuous measurements of the persistence of roadkill, and we expected to find that some animals would remain in place for only a few minutes or hours. We used this empirical distribution of carcass persistence times to simulate the roadkill removal process for a large number of carcasses. This simulation helped us investigate the potential impact of length-biased sampling on persistence estimates depending on the frequency of new roadkill recruitment. Subsequently, we conducted a second field experiment, this time using daily road surveys to recruit roadkill carcasses (similar to S. M. Santos et al., 2011). This allowed us to experimentally compare persistence estimates obtained from both experimental approaches, *i.e.,* carcass trials vs. recruiting roadkill with daily monitoring. Finally, we assessed the complexity of the roadkill persistence estimates in the study region by modelling the effects of several known predictors of roadkill persistence: traffic volume, rainfall, daily temperature, and species’ taxa and mean bodymass. As few studies have actually managed to record the mechanism by which roadkill is removed, we also leveraged the data provided by camera traps data to explore the causes of roadkill removal and their potential relation to roadkill persistence predictors, which may help explain observed roadkill persistence patterns.

## METHODS

All analyses were conducted using R version 4.4.1 (R Core Team, 2024).

### Simulation of roadkill persistence

We conducted a simulation of the appearance and removal of n=5000 animal roadkill on a roadway over a period of one month (31 days). Roadkill appearance was generated at random intervals following a Poisson distribution (Shoenfeld, 2004). To model the persistence of each simulated roadkill in the most realistic way possible, we used empirical data on persistence derived from a controlled experiment involving fresh carcasses intentionally placed on the road and monitored continuously with camera traps (see after, *Experiment setup,* n=73). We fitted several candidate distributions (exponential, log-normal, Weibull, gamma) to the observed data, following Huso (2011), and selected the distribution that best approximated the persistence patterns using the goodness-of-fit tests provided in the R package *fitdistrplus (Delignette-Muller & Dutang, 2015)*. The optimal distribution derived from this analysis was then used to generate a persistence duration for each simulated roadkill. For the purpose of simplification, we did not simulate any instances of censoring in the dataset, assuming that all carcasses were successfully monitored continuously from their placement on the road until their removal. This framework effectively simulated an experiment in which carcasses were deliberately placed on a roadway and monitored to evaluate their persistence.

In order to assess the impact of the recruitment frequency, we also devised a simulated monitoring protocol for roadkill. This protocol involved monitoring the road every 24 hours to identify new roadkill and track their presence every day until removal. For the sake of completeness, we also conducted simulations using monitoring frequencies of 12, 6 and 2 hours to demonstrate that the length bias decreases as the difference between carcass persistence and monitoring frequency is reduced. For each survey frequency, we assessed which of the simulated carcasses entered the study: not all were detected during these monitoring sessions, as some appeared and disappeared between scheduled monitoring events, and were therefore never recorded (Fig. 1). For the carcasses that were discovered, we recorded the number of consecutive monitoring sessions they remained present for, thereby allowing us to quantify their persistence as it would have been documented in a real experimental setting. We then compared the actual median persistence of the roadkill, and the measured persistence of the roadkill discovered during periodic road monitoring events using Kaplan-Meier survival curves.

**Figure 1:**
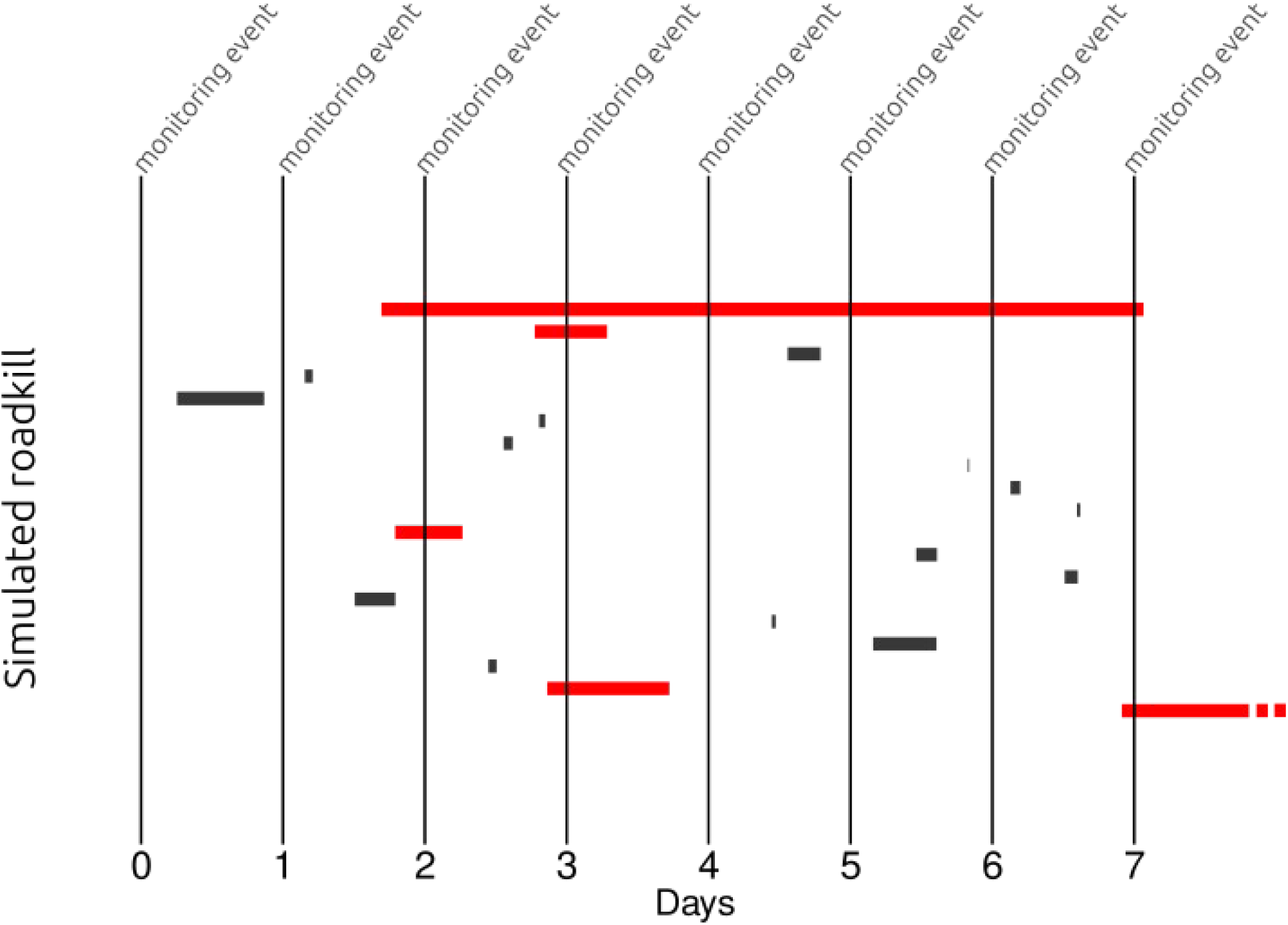
Simulation of an experiment to monitor roadkill persistence. Each animal appears on the roadway at a random time following a Poisson process and remains present for a specified duration, derived from a distribution fitted to empirical data collected from a controlled experiment. The experimental approach involves daily monitoring events, during which new carcasses are documented and the persistence of previously recorded roadkill is recorded. As the persistence of many simulated roadkill is shorter than the frequency of monitoring, only a limited number of carcasses (red) are present during the monitoring sessions and therefore recruited in the study.

### Experimental setup

We conducted the experiments in the Auvergne-Rhône-Alpes region of central-east France, between November and August. This region is characterized by a diverse landscape, dominated by artificial surfaces (41.6%) and forested or semi-natural landscapes (49.7%, Corine Land Cover, European Environment Agency, 2019). During the experimental sessions, average daily temperatures ranged between -1.5°C and 27°C, and average daily cumulated rainfall ranged between 0mm and 37.3mm. In both experimental setups, we considered that a carcass on the road was ‘present’ if it was visible from the drivers’ perspective, following Bénard et al. (2024), meaning that remains that were either too degraded, or had been placed somewhere there no longer visible (such as in the road ditch), were considered removed.

The first experimental approach measured the persistence of animal carcasses placed on the road by experimenters to simulate roadkill (thereafter: *intentionally placed* experiment). We randomly selected 15 roads in peri-urban or rural areas covering a range of traffic from 500 to 14 000 vehicles per day, representative of the diversity of traffic volumes for the road network of the area (Ministère de la Transition écologique, 2021). High-traffic roads (over 14 000 vehicles per day), specifically highways, were excluded for the observer’s safety (AB). We set up “Cuddeback 1279” camera-traps with a “Black Flash” module (night vision without a visible flash) at 30 different locations on the 15 roads (Fig. A1). We selected the exact locations randomly, while ensuring that traps could be reasonably hidden from the road to discourage theft. Traps located on the same road were placed at least 2km from one another. We placed each trap on a tree on the verge of the road, such that the entire width of the road was within the field of view of the camera, while also maintaining an approximately equal proportion of open (*e.g.,* agricultural fields) and closed (*e.g.,* forests) road-adjacent habitats. Traps were programmed to take 3 photographs when a movement triggered the sensors, plus an additional time-lapse photo every 3 hours regardless of movement. This time-lapse setting ensured that, should the trap not trigger as expected when the animal carcass disappeared from the road, we could estimate the persistence with a minimal accuracy of 3h. Camera traps also allowed us to determine the causes of disappearance, including being crushed by vehicle tires, removed by individuals, consumed by scavengers, and experiencing natural decay.

We procured 73 bird and mammal carcasses from a wildlife rehabilitation center, which were kept at -20°C and thawed 24h before the experiments. We selected animals that were not administered any medications while under the care of the rehabilitation center, in an effort to minimize the potential impact of the study on wildlife that may consume the carrion. We placed one animal at a time on the road surface in front of a camera trap, alternating between the center of the lane and the border between the road and shoulder. Note that we placed carcasses >5kg in weight exclusively on the shoulder due to safety concerns for drivers.

The second experimental approach measured roadkill persistence using daily road surveys to detect and follow animal carcasses (thereafter: *discovered roadkill* experiment). We recruited 12 volunteers through the LPO, a wildlife protection NGO, as well as via an announcement in a local newspaper. In addition, we enlisted the help of ecologists working for the *Syndicat de la rivière d’Ain Aval et de ses affluents,* a public syndicate that aims at protecting biodiversity. Each participant provided a set of roads they accepted to cover by car daily. They were tasked with detecting and monitoring roadkill carcasses present in their survey area. Volunteer participants sent a picture to the authors via phone or email every day, along with the location of the animal and a comment on the visibility of the animal from the road (whether it was visible to drivers or not).

Syndicate participants were instructed to also set up a camera trap on the road in order to monitor the disappearance of the discovered roadkill continuously (see above for camera trap settings). Both sets of participants were instructed not to displace roadkill detected during the experiment. In total, volunteers detected and monitored 55 road fatalities, and syndicate workers followed the disappearance of 9 animal carcasses with camera traps.

The 31 mammalian species used in the *intentionally placed* experiment ranged in weight from brown rat (*Rattus norvegicus*, 300g) to roe deer (*Capreolus capreolus*, 25kg), and the 42 birds from blue tit (*Cyanistes caeruleus*, 20g) to Eurasian eagle-owl (*Bubo bubo,* 2.5kg). Participants of the *discovered roadkill* experiment recorded the disappearance of 43 mammals, from brown rat (*Rattus norvegicus*, 300g) to roe deer (*Capreolus capreolus*, 25kg) and 20 bird from European robin (*Erithacus rubecula*, 20g) to common buzzard (*Buteo buteo,* 1.01kg). The *discovered roadkill* experiment tended to include heavier species (median bodymass: 1.3kg, Q1: 0.43kg, Q2: 5kg) compared to the experiment with animals intentionally placed (median bodymass: 0.72kg, Q1: 0.13kg, Q2: 1.0kg), possibly because small species are not easy to detect while driving. The daily mean temperature ranges and cumulative rainfall were comparable in both experiments, but the range of road traffic volumes was wider in the experimental approach with daily surveys (Fig. A2). The main difference between the experiments was the inclusion of 4 observations on highways—a category of roads typically heavily frequented, fenced, and inaccessible to pedestrians an—into the *discovered roadkill* experiment.

### Predictors of roadkill persistence

We retrieved the average body mass (continuous variable) of species present in both experiments from the various sources (Geptner et al., 1988; Hewison et al., 2002; Pettorelli et al., 2002; Sherfy et al., 2006; Herberstein et al., 2022; Tobias et al., 2022; Tranquillo et al., 2024). We retrieved the annual average volume of traffic (continuous variable) of the road from official sources (Conseil départemental de l’Ain, 2022; Ministère de la Transition écologique, 2021). The mean daily temperature (continuous variable) and daily cumulated rainfall (continuous variable) corresponded to the day the carcass was placed on the road (*intentionally placed* experiment) or the day the roadkill was discovered (*discovered roadkill* experiment) and were retrieved from public weather datasets (Larvor et al., 2020). All continuous predictors were scaled using R function *scale*. For both datasets, we considered participant ID or camera trap ID as random effects.

### Statistical analyses

We used Cox-Proportional Hazards models to estimate the median persistence of roadkill, and assess the influence of various predictors, using R package *survival* (Therneau, 2024). However, 39% of the camera traps did not activate upon the disappearance the animal during our *intentionally placed* experiment (either due to sensors error and battery failure). In consequence, some disappearances were only identified to have occurred within a known time interval sometimes spanning multiple days. Failure to account for interval-censoring in survival models can result in biased survival estimates in either direction and produce misleadingly low confidence intervals (Radke, 2003). To address this, we used survival models capable of incorporating interval-censoring through the R package *icenReg* (Anderson-Bergman, 2017) when necessary. Contrary to the *survival* package, *icenReg* does not support mixed models. For each model, we verified the proportional hazards assumption by plotting the Schoenfeld residuals (Schoenfeld, 1982) and the absence of multicollinearity by computing the variance inflation factors (R package *performance,* Lüdecke et al., 2021).

We first explored the impact of the experimental approach on the estimation of roadkill persistence by computing the Kaplan-Meier median persistence estimates (time for 50% of the carcasses to disappear) for both *intentionally placed* and *discovered roadkill*. We then modelled the effect of the experimental approach by analysing the datasets conjointly. We first degraded all data retrieved from camera traps as if the persistence had been measured daily instead, to ensure homogeneity across all methodologies (n=137, interval-censored model, no random effects). Next, we only used the camera trap data—the only method that could record persistence continuously—from both approaches (n=82, interval-censored model, no random effects). We included the species’ bodymass, road traffic volume, mean temperature and cumulated rainfall as co-variables to control for confounding factors and accurately capture the effect of the experimental method on recording roadkill persistence.

We also explored the relations between carcass persistence and preditors. Instead of relying on previous combined models, with all data from two inhomogeneous methodologies, we analysed the experiments separately to better account for the specificities of each dataset. In the *discovered roadkill* experiment, which did not contain interval-censoring, we were able to model the effects of traffic volume, species’ mean body mass, daily temperature, and cumulative rainfall, including participant ID as a random effect. In the *intentionally placed* experiment, where camera trap failures introduced several instances of interval-censoring, we modelled the effects of traffic volume, species’ mean body mass, daily temperature, and cumulative rainfall on roadkill persistence with no random effects.

Finally, we explored the importance of carcass removal mechanim in the persistence of roadkill: in all instances where the camera traps were able to capture the cause of roadkill disappearance, we tested whether the cause of removal from the road led to differences in persistence (interval-censored model, no random effects), and we applied a multinomial linear regression (R package *mclogit; Elff, 2022)*, including the camera trap number as a random effect, to evaluate whether the probability that roadkill removal was caused by removal by an individual, scavenging, decay or dismemberement by vehicles could be correlated to weather, road or species characteristics.

## RESULTS

### Simulation of roadkill disappearance

The empirical dataset of carcass persistence showed that 25% had disappeared within the first 2h13 (95% CI: 0h30, 5h19), 50% in 7h25 (95% CI: 3h54, 23h18) and 75% in 66h28 (95% CI: 18h, 138h51). The log-normal distribution fitted the empirical data better compared to the exponential, Weibull, and gamma distributions. Specifically, the log-normal distribution had a significantly lower Anderson-Darling (AD) statistic (AD: 0.205) compared to the exponential (AD: 33.783), Weibull (AD: 0.703), and gamma (AD: 2.739) distributions, indicating a closer alignment with the empirical data, particularly in the tails of the distributions (Engmann & Cousineau, 2011). Additionally, the log-normal model had a lower Akaike’s Information Criterion (AIC: 600.15) compared to the exponential (AIC: 721.70), Weibull (AIC: 611.11), and gamma (AIC: 632.06), further supporting its selection. The log-normal distribution, characterized by a heavy right tail, would tend to indicate that most roadkill was removed promptly, while some carcasses remained for extended periods. The persistence of roadkill was simulated using parameters log(*μ*) = 2.31 and *σ* = 2.04.

During the simulated daily monitoring, only 50.9% of the simulated carcasses were detected, while the remainder were present on the road exclusively between monitoring events and were therefore never recorded. The Kaplan-Meier estimates indicated that the true median survival time was approximately 9h58 (95% CI: 9h18,11h), while the median for the carcasses that were detected during the daily surveys was 38h18 (95% CI: 36h24,40h42), corresponding to an overestimation of the true carcass persistence by a factor 3.84 (Fig. 2). Performing monitoring sessions every 12 hours resulted in an overestimation by a factor of 2.70 (median persistence: 26h54, 95% CI: 25h12, 28h24), while monitoring the road every 6 hours reduced the overestimation to a factor of 2 (median persistence: 19h55, 95% CI: 18h55, 21h12), and monitoring every 2 hours, a factor of 1.41 (median persistence: 14h06, 95% CI: 13h12, 15h06), as more frequent surveys were able to detect carcasses more homogeneously (see Fig. A3 for the corresponding survival curves).

**Figure 2:**
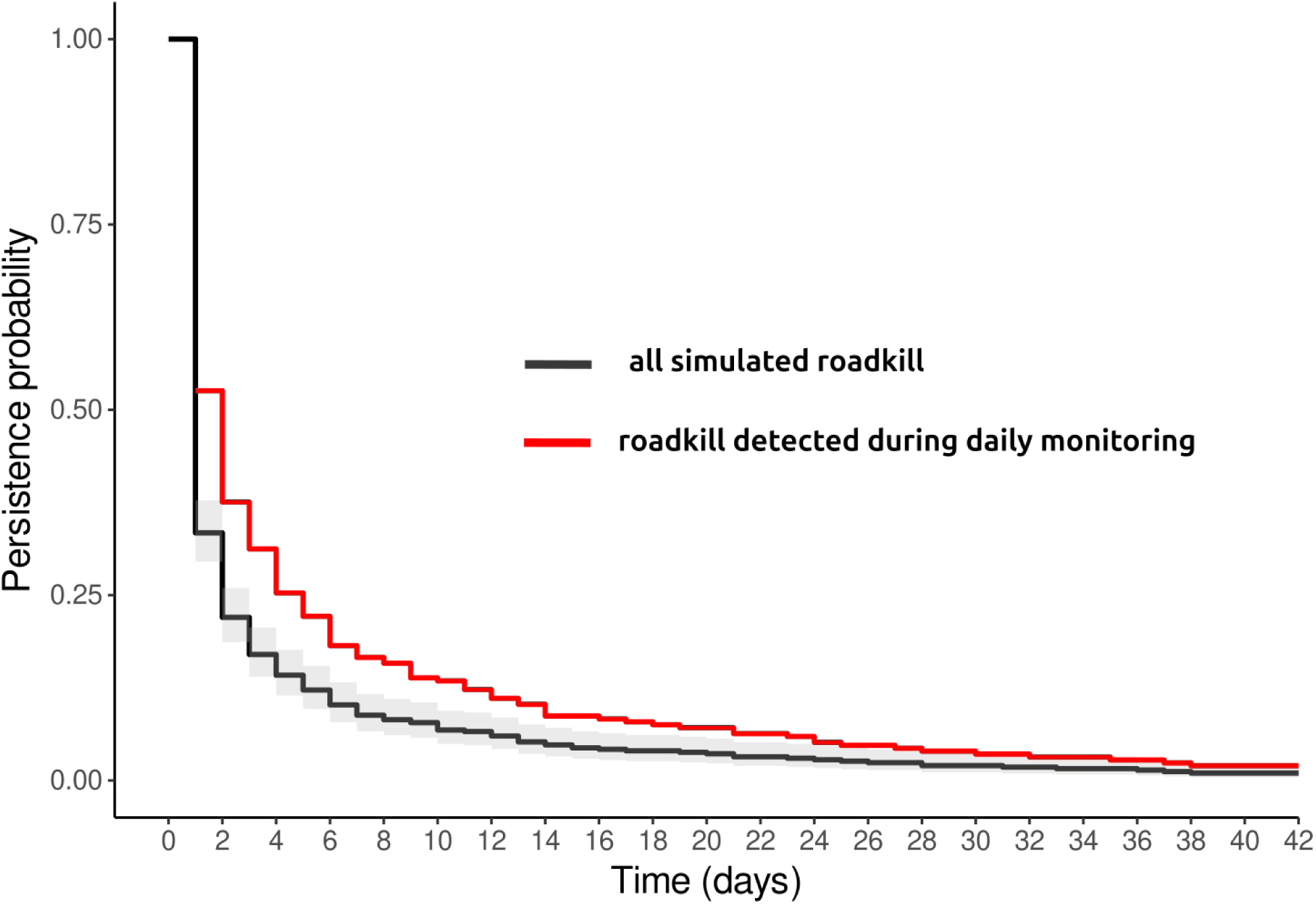
Kaplan-Meier survival curves of 5000 simulated animal roadkill. Daily monitoring of animal carcasses found on the road results in an overestimation of persistence due to a bias towards recording longer-lived carcasses over those with shorter persistence that falls in-between monitoring events.

### Experimental comparison of methodologies for recording roadkill persistence

The median persistence time of the animals placed on the road and monitored with camera traps was 7h25 (95% CI: 3h54, 23h18). Roadkill discovered and monitored daily persisted on average 2 days (95% CI: 1, 3). The median persistence of the 9 roadkills discovered and monitored with camera traps was 46h14 (95% CI: 20h30, NA^1^). The Kaplan-Meier survival curves for both *placed* and *discovered* are shown in figure 3.

**Figure 3:**
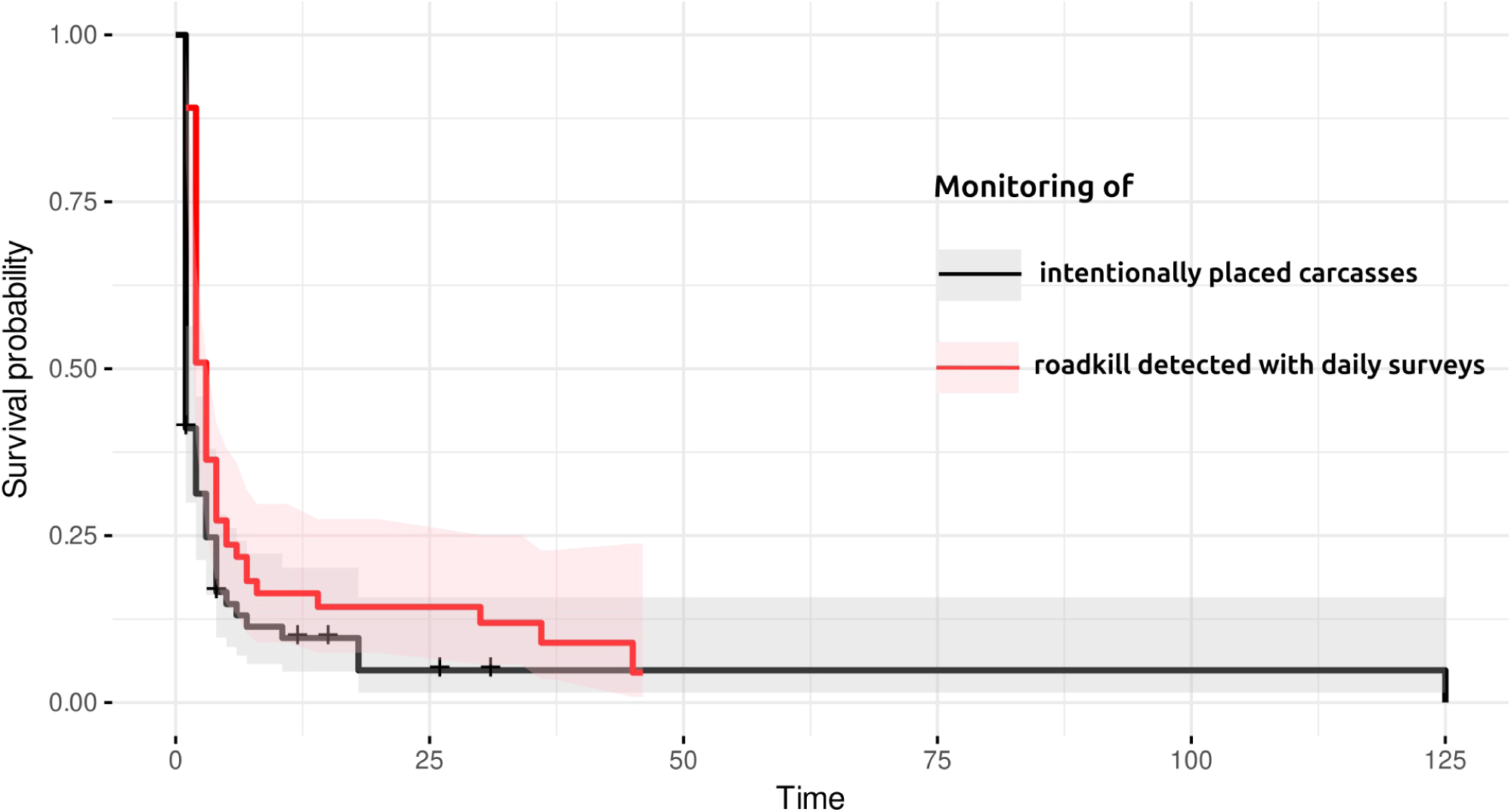
Kaplan-Meier survival curves of the persistence of animal carcasses on the road (hours). Two experimental approaches are tested: following the persistence of roadkill discovered during daily road surveys (red, n=64), and intentionally placing fresh animal carcasses on roads to monitor their disappearance (black, n=73).

However, we found no statistical evidence that the survival curves differed according to the experimental approach (Hazards Ratio=1.03 (95% CI: 0.65–1.65), p=0.887) after accounting for the other sources of confusion. In fact, the Cox-PH model for both datasets analysed conjointly showed no significant effects of the covariates: road traffic volume (Hazard Ratio=0.77 (95% CI: 0.59– 1.02), p=0.072), species’ mean body mass (HR=1.04 (95% CI: 0.79–1.37), p=0.802), average temperature (HR=1.16 (95% CI: 0.93–1.44), p=0.199), taxa (HR=0.79 (95% CI: 0.49–1.26), p=0.369). Similarly, when analysing only camera trap data (continuous monitoring only), we found no effect of the experimental approach (HR=1.41 (95% CI: 0.23–8.67), p=0.700) on roadkill persistence, nor of the other variables: road traffic volume (HR=0.96 (95% CI: 0.62–1.50), p=0.865), species’ mean body mass (HR=1.11 (95% CI: 0.64–1.91), p=0.720), average temperature (HR=1.03 (95% CI: 0.78–1.36), p=0.827), taxa (HR=0.62 (95% CI: 0.31–1.25), p=0.106).

### Effects of predictors on the speed and cause of roadkill disappearance

For the *intentionally placed* experiment, none of the predictors showed an effect on the rate of removal of roadkill (Table 1). However, when analysing the *discovered roadkill* experiment dataset separately, we find that lower volumes of road traffic and higher temperatures were associated with longer roadkill persistence. Through the use of camera traps, we were able to determine the cause of removal for 49 out of the 82 animals monitored. For the remaining observations, we sometimes encountered technical difficulties with the traps, including battery failures (n=3), misalignment of the traps with the placement of the animal carcass on the road (n=3), theft of the trap (n=2), and a trap malfunction (n=1). However, in 20 cases the trap simply failed to activate during the animal’s removal, and its disappearance was only evident from the time-lapse photographs that were captured every 3 hours. We find that animal carcasses were mainly displaced by passers-by on foot, bike, or in cars (stopping the vehicle to displace the animal, 55.8% of cases). In most instances, the carcass was still present but no longer visible from the road (placed in a ditch, dragged several meters away), although 1 red fox *Vulpes vulpes* was apparently collected. Participants of the *discovered* roadkill also reported finding carcasses hidden in road ditches. Other animals were repeatedly crushed by vehicles until the remains were no longer identifiable (41.8% of cases). Only one common swift *(Apus apus*) and one European rabbit (*Oryctolagus cuniculus)* were seen consumed by scavengers (identified carrionr-feeders: European badger *Meles meles,* common buzzard *Buteo buteo,* and red fox *Vulpes vulpes*). Therefore, only two causes of roadkill removal could be used in the subsequent analyses: “removed by someone” and “crushed by vehicles”.

Animals removed by members of the public lasted an average of 4h44 (95% CI: 2h14, 6h07), while animals removed through repeated crushing lasted an average of 20h28 (95% CI: 10h41, 82h58). The Cox-PH model showed that this difference in carcass persistence was statistically significant (Hazard Ratio=0.47 (95% CI: 0.27–0.84), p=0.004). The logistic regression showed no significant correlation between the cause of animal removal and the volume of traffic on the road (Odds Ratio=2.52 (95% CI: 0.92–6.93), p=0.078), the cumulative rainfall (OR=1.10 (95% CI: 0.38–3.18), p=0.823), or the average daily temperature (OR=0.95 (95% CI: 0.37–2.43), p=0.904). However, heavier species had higher odds of being removed by people than they had of being crushed by vehicles (OR=8.00e-7 (95% CI: 2.23e-10–2.87e-2), p=0.0179), and there was marginal evidence for mammals being more often removed by people (OR=12.40 (95% CI: 0.94–162.90), p=0.0566).

## DISCUSSION

Information on roadkill persistence is fundamental for conducting research on wildlife-vehicle collisions. In this study, the average persistence for a carcass placed on a road was approximately 8 hours, consistent with prior studies conducted on a variety of roadkill species and road conditions, where most roadkill disappeared within the first 24h (Beckmann & Shine, 2015; Bénard et al., 2024; Brzeziński et al., 2012; Ratton et al., 2014; Santos et al., 2016; Stewart, 1971). Such short persistence times raise concerns for the quality of roadkill data, as monitoring schemes with a frequency of >1 day are likely to miss an important proportion of the roadkill (Henry et al., 2021). Our results suggest that even surveys performed twice a day (with an interval of 12h) will miss 58% of the mortality on roads, while daily surveys will miss 69% of the mortality. While collisions with vehicles are not currently recognized as one of the primary human-induced causes of mortality in terrestrial animals (Hill et al., 2019), our findings suggest that this may be due to a significant underestimation of the actual impact of roadways on wildlife. However, increasing monitoring frequencies to accurately record road mortality may not always be a realistic answer, as costs may not be sustainable (Henry et al., 2021). Ad-hoc methods for roadkill data correction, using estimates of the persistence, may represent more viable solutions (Teixeira et al., 2013).

The exponential distribution was not the best fit for the carcass persistence data obtained from the carcass trial experiments; indicating that the hazard rate for roadkill removal is not constant over time (Bender et al., 2005). In practice, this indicates a rapid decline in the proportion of remaining carcasses at the outset, which subsequently slows down: animal roadkill will initially vanish quickly, and for carcasses that are still present on roads after a few days, the probability of persistence increases. The removal of roadkill is frequently attributed to scavengers activity (Antworth et al., 2005; Boves & Belthoff, 2012; Hubbard & Chalfoun, 2012; Slater, 2002). Consequently, the effects of predictors are often interpreted through this lens: the extended persistence of carcasses on highly frequented roads is explained as a result of the barrier effect that these roads impose on the presence of scavengers (R. A. L. Santos et al., 2016; S. M. Santos et al., 2011). When models indicate that larger species remain in place longer, authors conclude that this phenomenon is due to smaller carrion being more susceptible to being carried away by scavengers (S. M. Santos et al., 2011). Furthermore, variations in persistence depending on the time of day, season, and weather can be explained by variations in the activity and composition of the local scavenger guilds (Antworth et al., 2005; Beckmann & Shine, 2015; Guinard et al., 2012). Following this rationale, one could infer that carcasses are predominantly appealing to scavengers while fresh, resulting in elevated rates of removal initially (Guinard et al., 2012; Henry et al., 2021). Meanwhile, the few carcasses that remain will begin to decompose or harden, rendering them unlikely to be consumed (S. M. Santos et al., 2011; Winton et al., 2018).

However, camera trap data indicated minimal involvement of scavengers in roadkill removal in the central-east France, where various carrion feeders are present (Faune France, 2024). Experiments were conducted in late fall, winter, and early spring, seasons when scavenging could be expected to increase due to food scarcity (Prosser et al., 2008; Selva et al., 2005). Baiting roads may have led to carcasses that didn’t align with regular carrion hotspots expected by scavengers (Bracey et al., 2016). Additionally, traps failed to trigger in 29% of cases upon carcass disappearance, suggesting potential underreporting of scavenger activity, a limitation also highlighted by Winton et al. (2018). We observe that at least one-third of carcasses are removed by passing vehicles, and another third are continuously crushed until the remains are no longer detectable during surveys. Removal by humans primarily targeted larger animals, possibly due to their increased visibility or the perception that they pose a security risk to other drivers or potential scavengers. Some researchers have suggested that roadkill may hold symbolic significance for the public, representing the destruction of nature or evoking feelings of guilt (Lunney, 2013). This emotional response could lead road users to remove carcasses in an effort to conceal road mortality or as a form of mourning (Desmond, 2013; Lulka, 2008). Human road clean-up and dismemberment by vehicles may also explain why hazard rates were not constant over time: carcasses located directly in the path of vehicle tires may disappear quickly (Bénard et al., 2024; R. A. L. Santos & Ascensão, 2019), resulting in an initial rapid decrease in carcass numbers that slows down over time as only unreachable carcasses remain. Additionally, people might promptly remove carcasses perceived as immediate safety risks or if they are near convenient parking spots. Once carcasses start decaying or become disfigured, people may be reluctant to displace them. Previous studies have discussed that some carcasses will remain in place for extended periods of time (S. M. Santos et al., 2011).

Our simulations indicate that sampling roadkill carcasses with regular road surveys may lead to overestimating the persistence. This is due to under-sampling short-lived carcasses, which are removed from roads more quickly than the survey intervals. Length-biased sampling is a well-recognized issue in the literature (Shen et al., 2017), especially in medical studies (e.g., cancer research, (Kay & Witte, 1991). However, it should be noted that the simulation relies on fitting a survival curve to empirical data obtain through *carcass trials* to represent true carcass persistence patterns. Although this has never been formally demonstrated, carcass trials may not precisely replicate roadkill patterns as encountered by local carrion-feeders (Bracey et al., 2016). This mismatch could distort the distribution of persistence time. Our simulation also employed a Poisson distribution to model the timing of collisions, given the limited research available that records the precise timing of collisions. Therefore, while we suspect that that length bias is present in roadkill persistence studies via periodic monitoring; however, in the absence of an unbiased reference for true survival distributions, its exact magnitude and nature remain unquantifiable.

Experimental results reflected the trend obtained through simulation, with median persistence estimates from our daily recruitment surveys (*discovered roadkill)* being longer than those from trials using carcasses placed on roads (*intentionally placed),* and mean survival curves being distinct (Fig. 3). We were however unable to demonstrate a statistically significant difference between the survival curves from the two experiments after controlling for all the sources of variation (animal body mass, taxa, road trafic, and weather). This may be due to insufficient statistical power, as obtaining large sample sizes was challenging: we depended on recruiting volunteers who were able to monitor roadways daily for extended durations, in addition to facing limitations regarding the availability of fresh, unmedicated carcasses for the carcass trials.

The variations in roadkill persistence estimates across studies (Table 1) can be attributed to differences in species composition, road characteristics, weather patterns, and the presence of scavenger guilds in the study area (Antworth et al., 2005; S. M. Santos et al., 2011). Additionally, we show that the cultural tendency to remove roadkill may significantly impact the speed of roadkill removal. Several authors have cautioned that roadkill persistence estimates, which are crucial for adjusting roadkill rates, should be generalized between studies with caution (Brzeziński et al., 2012; S. M. Santos et al., 2011). In central-east France, we found little evidence of the spatio-temporal patterns usually present in roadkill persistence, in contradiction to previous European studies (Bénard et al., 2024; Brzeziński et al., 2012; Guinard et al., 2012; S. M. Santos et al., 2011; Slater, 2002), although several studies show that these patterns are not always consistent (Antworth et al., 2005; Henry et al., 2021). In fact, we find not emprical support for different in roadkill disappearance speed depending on species and rainfall. We find limited support for an effect of temperature and road traffic, in accordance with Santos et al. (2011). These disparities in the effect of covariates between the experimental methodologies may reflect the greater variability in sampled locations in the *discovered* experiment (which was not artificially limited to less frequented roads).

## CONCLUSION

Roadkill monitoring is a widely used method for assessing the impact of vehicular traffic on wildlife. However, this approach is subject to potential biases due to the relatively brief persistence of roadkill, which may be removed by individuals, scavengers, or vehicles. This bias can affect the accurate identification of dangerous road segments and species at risk, as well as potentially leading to an underestimation of biodiversity loss attributable to road mortality. Despite this issue, there have been relatively few studies focused on estimating roadkill persistence. Our findings corroborate previous research, indicating that roadkill typically disappears within a matter of hours. Therefore, even daily monitoring could introduce biases in roadkill counts. Additionally, it appears that the rate at which roadkill disappears may vary according to the duration for which the animal has already persisted, resulting in the potential for length-biased sampling when estimating persistence through regular road monitoring. Accurately describing the distribution of roadkill persistence times may therefore present a greater challenge than previously anticipated.

## Footnotes

1 Failure to compute the upper bound of the confidence interval is due to the small sample size and right-censoring. Note that, when excluding observations conducted on highways from the *discovered* dataset, the median persistence of roadkill remained at 2 days (95% CI: 1, 3).

## REFERENCES

Anderson-Bergman, C. (2017). Bayesian Regression Models for Interval-censored Data in R. The R Journal, 9(2), 487. 10.32614/RJ-2017-050

Antworth, R. L., Pike, D. A., & Stevens, E. E. (2005). Hit and run : Effects of scavenging on estimates of roadkilled vertebrates. Southeastern Naturalist, 4(4), 647–656. 10.1656/1528-7092(2005)004[0647:HAREOS]2.0.CO;2

Aresco, M. J. (2003). Highway mortality of turtles and other herpetofauna at Lake Jackson, Florida, USA, and the efficacy of a temporary fence/culvert system to reduce roadkills. https://escholarship.org/uc/item/0kr0x064

Barrientos, R., Martins, R. C., Ascensão, F., D’Amico, M., Moreira, F., & Borda-de-Água, L. (2018). A review of searcher efficiency and carcass persistence in infrastructure-driven mortality assessment studies. Biological Conservation, 222(November 2017), 146–153. 10.1016/j.biocon.2018.04.014

Beckmann, C., & Shine, R. (2015). Do the numbers and locations of road-killed anuran carcasses accurately reflect impacts of vehicular traffic? The Journal of Wildlife Management, 79(1), 92–101. 10.1002/jwmg.806

Bénard, A., Bonenfant, C., & Lengagne, T. (2024). Traffic and weather influence on small wildlife carcass persistence time on roads. Transportation Research Part D: Transport and Environment, 126, 104012. 10.1016/j.trd.2023.104012

Bender, R., Augustin, T., & Blettner, M. (2005). Generating survival times to simulate Cox proportional hazards models. Statistics in Medicine, 24(11), 1713–1723. 10.1002/sim.2059

Boves, T. J., & Belthoff, J. R. (2012). Roadway mortality of barn owls in Idaho, USA. The Journal of Wildlife Management, 76(7), 1381–1392. 10.1002/jwmg.378

Bracey, A. M., Etterson, M. A., Niemi, G. J., & Green, R. F. (2016). Variation in bird-window collision mortality and scavenging rates within an urban landscape. The Wilson Journal of Ornithology, 128(2), 355–367. 10.1676/wils-128-02-355-367.1

Brzeziński, M., Eliava, G., & Żmihorski, M. (2012). Road mortality of pond-breeding amphibians during spring migrations in the Mazurian Lakeland, NE Poland. European Journal of Wildlife Research, 58(4), 685–693. 10.1007/s10344-012-0618-2

Cabrera-Casas, L. X., Robayo-Palacio, L. M., & Vargas-Salinas, F. (2020). Persistence of snake carcasses on roads and its potential effect on estimating roadkills in a megadiverse country. Amphib. Reptile Conserv., 14(1).

Collinson, W. J., Parker, D. M., Bernard, R. T. F., Reilly, B. K., & Davies-Mostert, H. T. (2014). Wildlife road traffic accidents : A standardized protocol for counting flattened fauna. Ecology and Evolution, 4(15), 3060–3071. 10.1002/ece3.1097

Conseil départemental de l’Ain. (2022). *Trafic routier—Bilan* 2022 [Jeu de données]. https://www.ain.fr/app/uploads/2023/03/livret-comptages-routiers-2022-bilan-des-trafics-et-circulation-ain.pdf

Delignette-Muller, M. L., & Dutang, C. (2015). fitdistrplus : An R Package for Fitting Distributions. Journal of Statistical Software, 64, 1–34. 10.18637/jss.v064.i04

Desmond, J. (2013). Requiem for Roadkill : Death and Denial on America’s Roads. In Environmental Anthropology. Routledge.

Díaz, S., Settele, J., Brondízio, E. S., Ngo, H. T., Agard, J., Arneth, A., Balvanera, P., Brauman, K. A., Butchart, S. H. M., Chan, K. M. A., Garibaldi, L. A., Ichii, K., Liu, J., Subramanian, S. M., Midgley, G. F., Miloslavich, P., Molnár, Z., Obura, D., Pfaff, A., … Zayas, C. N. (2019). Pervasive human-driven decline of life on Earth points to the need for transformative change. Science, 366(6471), eaax3100. 10.1126/science.aax3100

Elff, M. (2022). mclogit : Multinomial Logit Models, with or without Random Effects or Overdispersion. https://CRAN.R-project.org/package=mclogit

Engmann, S., & Cousineau, D. (2011). Comparing distributions : The two-sample Anderson-Darling test as an alternative to the Kolmogorov-Smirnoff test. Journal of Applied Quantitative Methods, 6(3). http://jaqm.ro/issues/volume-6,issue-3/pdfs/jaqm_vol6_issue3.pdf#page=5

Etterson, M. A. (2013). Hidden Markov models for estimating animal mortality from anthropogenic hazards. Ecological Applications, 23(8), 1915–1925. 10.1890/12-1166.1

European Environment Agency. (2019). *CORINE Land Cover 2018 (vector), Europe*, *6-yearly— Version 2020_20u1, May* 2020 (Version 20.01) [FGeo,Spatialite]. European Environment Agency. 10.2909/71C95A07-E296-44FC-B22B-415F42ACFDF0

European Environment Agency. (2024). *Greenhouse gas emissions by aggregated sector* [Data Visualization]. https://www.eea.europa.eu/data-and-maps/daviz/ghg-emissions-by-aggregated-sector-5#tab-dashboard-02

Forman, R. T. T., & Alexander, L. E. (1998). Roads and their major ecological effects. Annual Review of Ecology and Systematics, 29(1998), 207–231. 10.1146/annurev.ecolsys.29.1.207

Franceschi, I. C., Gonçalves, L. O., Kindel, A., & Trigo, T. C. (2021). Mammalian fatalities on roads : How sampling errors affect road prioritization and dominant species influence spatiotemporal patterns. European Journal of Wildlife Research, 67(6), 97. 10.1007/s10344-021-01540-z

Gaston, K. J., & Holt, L. A. (2018). Nature, extent and ecological implications of night-time light from road vehicles. Journal of Applied Ecology, 55(5), 2296–2307. 10.1111/1365-2664.13157

Geptner, V. G. (Vladimir G., Nasimovich, A. A., Bannikov, A. G., & Hoffmann, R. S. (avec Smithsonian Libraries). (1988). Mammals of the Soviet Union. Washington, D.C. : Smithsonian Institution Libraries and National Science Foundation. http://archive.org/details/mammalsofsov212001gept

Gerow, K., Kline, N., Swann, D., & Pokorny, M. (2010). Estimating annual vertebrate mortality on roads at Saguaro National Park, Arizona. Human-Wildlife Interactions, 4(2), 283–292.

Girardet, X., Conruyt-Rogeon, G., & Foltête, J.-C. (2015). Does regional landscape connectivity influence the location of roe deer roadkill hotspots? European Journal of Wildlife Research, 61(5), 731–742. 10.1007/s10344-015-0950-4

Gomes, L., Grilo, C., Silva, C., & Mira, A. (2009). Identification methods and deterministic factors of owl roadkill hotspot locations in Mediterranean landscapes. Ecological Research, 24(2), 355–370. 10.1007/s11284-008-0515-z

Grilo, C., Koroleva, E., Andrášik, R., Bíl, M., & González-Suárez, M. (2020). Roadkill risk and population vulnerability in European birds and mammals. Frontiers in Ecology and the Environment, 18(6), 323–328. 10.1002/fee.2216

Guinard, E., Billon, L., Bretaud, J.-F., Chevallier, L., Sordello, R., & Witté, I. (2023). Comparing the effectiveness of two roadkill survey methods on roads. Transportation Research Part D: Transport and Environment, 121, 103829. 10.1016/j.trd.2023.103829

Guinard, E., Julliard, R., & Barbraud, C. (2012). Motorways and bird traffic casualties : Carcasses surveys and scavenging bias. Biological Conservation, 147(1), 40–51. 10.1016/j.biocon.2012.01.019

Henry, D. A. W., Collinson-Jonker, W. J., Davies-Mostert, H. T., Nicholson, S. K., Roxburgh, L., & Parker, D. M. (2021). Optimising the cost of roadkill surveys based on an analysis of carcass persistence. Journal of Environmental Management, 291, 112664. 10.1016/j.jenvman.2021.112664

Herberstein, M. E., McLean, D. J., Lowe, E., Wolff, J. O., Khan, M. K., Smith, K., Allen, A. P., Bulbert, M., Buzatto, B. A., Eldridge, M. D. B., Falster, D., Fernandez Winzer, L., Griffith, S. C., Madin, J. S., Narendra, A., Westoby, M., Whiting, M. J., Wright, I. J., & Carthey, A. J. R. (2022). AnimalTraits—A curated animal trait database for body mass, metabolic rate and brain size. Scientific Data, 9(1), 265. 10.1038/s41597-022-01364-9

Hewison, A. J. M., Gaillard, J. M., Angibault, J. M., Van Laere, G., & Vincent, J. P. (2002). The influence of density on post-weaning growth in roe deer Capreolus capreolus fawns. Journal of Zoology, 257(3), 303–309. 10.1017/S0952836902000900

Hill, J. E., DeVault, T. L., & Belant, J. L. (2019). Cause-specific mortality of the world’s terrestrial vertebrates. Global Ecology and Biogeography, 28(5), 680–689. 10.1111/geb.12881

Holderegger, R., & Di Giulio, M. (2010). The genetic effects of roads : A review of empirical evidence. Basic and Applied Ecology, 11(6), 522–531. 10.1016/j.baae.2010.06.006

Hubbard, K. A., & Chalfoun, A. D. (2012). An experimental evaluation of potential scavenger effects on snake road mortality detections. Herpetological Conservation and Biology, 7(2), 150–156.

Huso, M. M. P. (2011). An estimator of wildlife fatality from observed carcasses. Environmetrics, 22(3), 318–329. 10.1002/env.1052

Hwang, H.-M., Fiala, M. J., Park, D., & Wade, T. L. (2016). Review of pollutants in urban road dust and stormwater runoff : Part 1. Heavy metals released from vehicles. International Journal of Urban Sciences, 20(3), 334–360. 10.1080/12265934.2016.1193041

Kay, B. R., & Witte, D. L. (1991). The Impact of Cancer Biology, Lead Time Bias, and Length Bias in the Debate About Cancer Screening Tests. 23(2).

Langen, T. A., Machniak, A., Crowe, E. K., Mangan, C., Marker, D. F., Liddle, N., & Roden, B. (2007). Methodologies for Surveying Herpetofauna Mortality on Rural Highways. The Journal of Wildlife Management, 71(4), 1361–1368. 10.2193/2006-385

Larvor, G., Berthomier, L., Chabot, V., Le Pape, B., Pradel, B., & Perez, L. (2020). *MeteoNet, an open reference weather dataset by METEO FRANCE* [Jeu de données]. https://meteonet.umr-cnrm.fr/

Lüdecke, D., Ben-Shachar, M. S., Patil, I., Waggoner, P., & Makowski, D. (2021). performance : An R Package for Assessment, Comparison and Testing of Statistical Models. Journal of Open Source Software, 6(60), 3139. 10.21105/joss.03139

Lulka, D. (2008). The intimate hybridity of roadkill : A Beckettian view of dismay and persistence. *Emotion*, Space and Society, 1(1), 38–47. 10.1016/j.emospa.2008.08.008

Lunney, D. (2013). Wildlife roadkill : Illuminating and overcoming a blind spot in public perception. Pacific Conservation Biology, 19(4), 233–249. 10.1071/pc130233

Menger, T., Kindel, A., & Brack, I. V. (2023). Estimating roadkill rates while accounting for carcass detection and persistence using open-population capture–recapture models. Wildlife Research. 10.1071/WR22132

Ministère de la Transition écologique. (2021). *Trafic moyen journalier annuel sur le réseau routier national* [Jeu de données]. https://www.data.gouv.fr/fr/datasets/trafic-moyen-journalier-annuel-sur-le-reseau-routier-national/#/information

Moore, L. J., Petrovan, S. O., Bates, A. J., Hicks, H. L., Baker, P. J., Perkins, S. E., & Yarnell, R. W. (2023). Demographic effects of road mortality on mammalian populations : A systematic review. Biological Reviews. 10.1111/brv.12942

Nielsen, C., & Lang, R. S. (1999). Principles of screening. The Medical Clinics of North America, 83(6), 1323–1337, v. 10.1016/s0025-7125(05)70169-3

Ogletree, K. A., & Mead, A. J. (2020). What Roadkills Did We Miss in a Driving SurveyA Comparison of Driving and Walking Surveys in Baldwin County, Georgia.

Paraguassu-Chaves, C. A., Izidorio, A. de O., da Silva Júnior, N. P., Filho, A. K. D. B., Pereira, L. S., de Almeida, F. M., Neto, J. V. F., Barata, C. da S., Cavalcante, F. R. C., & Casara, H. N. (2020). Monitoring of Wildlife Mortality on a State Road in Rondônia, Western Amazon. International Journal of Advanced Engineering Research and Science, 7(8), 208–221. 10.22161/ijaers.78.21

Parris, K. M., & Schneider, A. (2009). Impacts of Traffic Noise and Traffic Volume on Birds of Roadside Habitats. Ecology and Society, 14(1). https://www.jstor.org/stable/26268029

Péron, G., Hines, J. E., Nichols, J. D., Kendall, W. L., Peters, K. A., & Mizrahi, D. S. (2013). Estimation of bird and bat mortality at wind-power farms with superpopulation models. Journal of Applied Ecology, 50(4), 902–911.

Pettorelli, N., Gaillard, J.-M., Van Laere, G., Duncan, P., Kjellander, P., Liberg, O., Delorme, D., & Maillard, D. (2002). Variations in adult body mass in roe deer : The effects of population density at birth and of habitat quality. Proceedings of the Royal Society of London. Series B: Biological Sciences, 269(1492), 747–753. 10.1098/rspb.2001.1791

Prosser, P., Nattrass, C., & Prosser, C. (2008). Rate of removal of bird carcasses in arable farmland by predators and scavengers. Ecotoxicology and Environmental Safety, 71(2), 601–608. 10.1016/j.ecoenv.2007.10.013

R Core Team. (2024). R: A Language and Environment for Statistical Computing. R Foundation for Statistical Computing. https://www.R-project.org/

Radke, B. R. (2003). A demonstration of interval-censored survival analysis. Preventive Veterinary Medicine, 59(4), 241–256. 10.1016/s0167-5877(03)00103-x

Ratton, P., Secco, H., & da Rosa, C. A. (2014). Carcass permanency time and its implications to the roadkill data. European Journal of Wildlife Research, 60(3), 543–546. 10.1007/s10344-014-0798-z

Rytwinski, T., Soanes, K., Jaeger, J. A. G., Fahrig, L., Findlay, C. S., Houlahan, J., Van Ree, R. D., & Van Der Grift, E. A. (2016). How effective is road mitigation at reducing road-killA meta-analysis. PLoS ONE, 11(11), 1–25. 10.1371/journal.pone.0166941

Santos, R. A. L., & Ascensão, F. (2019). Assessing the effects of road type and position on the road on small mammal carcass persistence time. European Journal of Wildlife Research, 65(1). 10.1007/s10344-018-1246-2

Santos, R. A. L., Santos, S. M., Santos-Reis, M., Picanço de Figueiredo, A., Bager, A., Aguiar, L. M. S., & Ascensão, F. (2016). Carcass Persistence and Detectability : Reducing the Uncertainty Surrounding Wildlife-Vehicle Collision Surveys. PLOS ONE, 11(11), e0165608. 10.1371/journal.pone.0165608

Santos, S. M., Carvalho, F., & Mira, A. (2011). How long do the dead survive on the roadCarcass persistence probability and implications for road-kill monitoring surveys. PLoS ONE, 6(9). 10.1371/journal.pone.0025383

Santos, S. M., Marques, J. T., Lourenço, A., Medinas, D., Barbosa, A. M., Beja, P., & Mira, A. (2015). Sampling effects on the identification of roadkill hotspots : Implications for survey design. Journal of Environmental Management, 162, 87–95. 10.1016/j.jenvman.2015.07.037

Schoenfeld, D. (1982). Partial residuals for the proportional hazards regression model. Biometrika, 69(1), 239–241. 10.1093/biomet/69.1.239

Seiler, A. (2001). Ecological Effects of Roads : A review. Uppsala: Swedish University of Agricultural Sciences.

Selva, N., Jędrzejewska, B., Jędrzejewski, W., & Wajrak, A. (2005). Factors affecting carcass use by a guild of scavengers in European temperate woodland. Canadian Journal of Zoology, 83(12), 1590–1601. 10.1139/z05-158

Shen, Y., Ning, J., & Qin, J. (2017). Nonparametric and Semiparametric Regression Estimation for Length-biased Survival Data. Lifetime data analysis, 23(1), 3–24. 10.1007/s10985-016-9367-y

Sherfy, M. H., Mollett, T. A., McGOWAN, K. R., & Daugherty, S. L. (2006). A Reexamination of Age-Related Variation in Body Weight and Morphometry of Maryland Nutria. Journal of Wildlife Management, 70(4), 1132–1141. 10.2193/0022-541X(2006)70[1132:AROAVI]2.0.CO;2

Shoenfeld, P. (2004). Suggestions regarding Avian Mortality Extrapolation. Prepared for the Mountaineer Wind Energy Center Technical Review Committee. https://docplayer.net/104547431-Suggestions-regarding-avian-mortality-extrapolation.html

Slater, F. M. (2002). An assessment of wildlife road casualties—The potential discrepancy between numbers counted and numbers killed. Web Ecology, 3, 33–42. 10.5194/we-3-33-2002

Stewart, P. A. (1971). Persistence of Remains of Birds Killed on Motor Highways. The Wilson Bulletin, 83(2), 203–204.

Taylor, B. D., & Goldingay, R. L. (2004). Wildlife road-kills on three major roads in north-eastern New South Wales. Wildlife Research, 31(1), 83–91. 10.1071/wr01110

Teixeira, F. Z., Coelho, A. V. P., Esperandio, I. B., & Kindel, A. (2013). Vertebrate road mortality estimates : Effects of sampling methods and carcass removal. Biological Conservation, 157, 317–323. 10.1016/j.biocon.2012.09.006

Teixeira, F. Z., Coelho, I. P., Esperandio, I. B., Rosa Oliveira, N., Porto Peter, F., Dornelles, S. S., Rosa Delazeri, N., Tavares, M., Borges Martins, M., & Kindel, A. (2013). Are road-kill hotspots coincident among different vertebrate groups? Oecologia Australis, 17(1), 36–47. 10.4257/oeco.2013.1701.04

Therneau, T. (s. d.). A package for survival analysis in R.

Tobias, J. A., Sheard, C., Pigot, A. L., Devenish, A. J. M., Yang, J., Sayol, F., Neate-Clegg, M. H. C., Alioravainen, N., Weeks, T. L., Barber, R. A., Walkden, P. A., MacGregor, H. E. A., Jones, S. E. I., Vincent, C., Phillips, A. G., Marples, N. M., Montaño-Centellas, F. A., Leandro-Silva, V., Claramunt, S., … Schleuning, M. (2022). AVONET : Morphological, ecological and geographical data for all birds. Ecology Letters, 25(3), 581–597. 10.1111/ele.13898

Tranquillo, C., Wauters, L. A., Santicchia, F., Panzeri, M., Preatoni, D., Martinoli, A., & Bisi, F. (2024). The advantage of living in the city : Effects of urbanization on body size and mass of native and alien squirrels. Urban Ecosystems, 27(1), 51–61. 10.1007/s11252-023-01435-8

Winton, S. A., Taylor, R., Bishop, C. A., & Larsen, K. W. (2018). Estimating actual versus detected road mortality rates for a northern viper. Global Ecology and Conservation, 16, e00476. 10.1016/j.gecco.2018.e00476

