## Appendix for "Experimental and Simulated Approaches to Roadkill Persistence: Implications for Road Mortality Assessment"

SUPPLEMENTARY MATERIAL

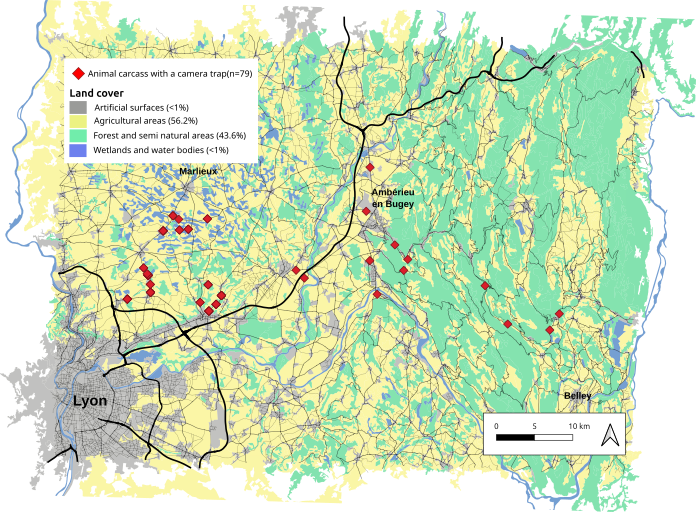

Figure A1: Map of the area in which we conducted the carcass persistence experiments for the “intentionally placed” approach. Each red point represents an animal carcass we placed on the road surface or shoulder, with an infra-red camera trap installed in the road side vegetation such that the animal is visible on the photographs. We used a total of 26 species donated by a wildlife rehabilitation center.

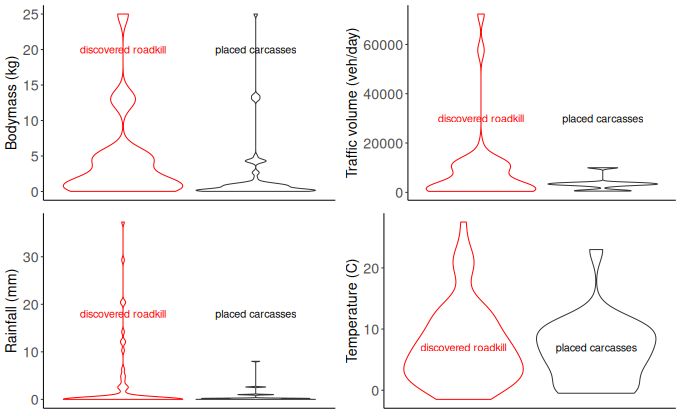

Figure A2: Comparison of the (a) mean species’ bodymass, (b) annual average daily volume of traffic of the road where the carcass was located, (c) daily cumulated rainfall of the day of the discovery or placement of carcasses and (d) daily average temperature of the day of the discovery or placement of carcasses. In the “discovered roadkill” approach (red), roadkill was detected with daily surveys, while in the “placed carcasses” approach (black) we deposited animal carcasses on the road to simulate roadkill. Roads with high volumes of traffic were intentionally not included in the “placed” approach due to safety concerns.

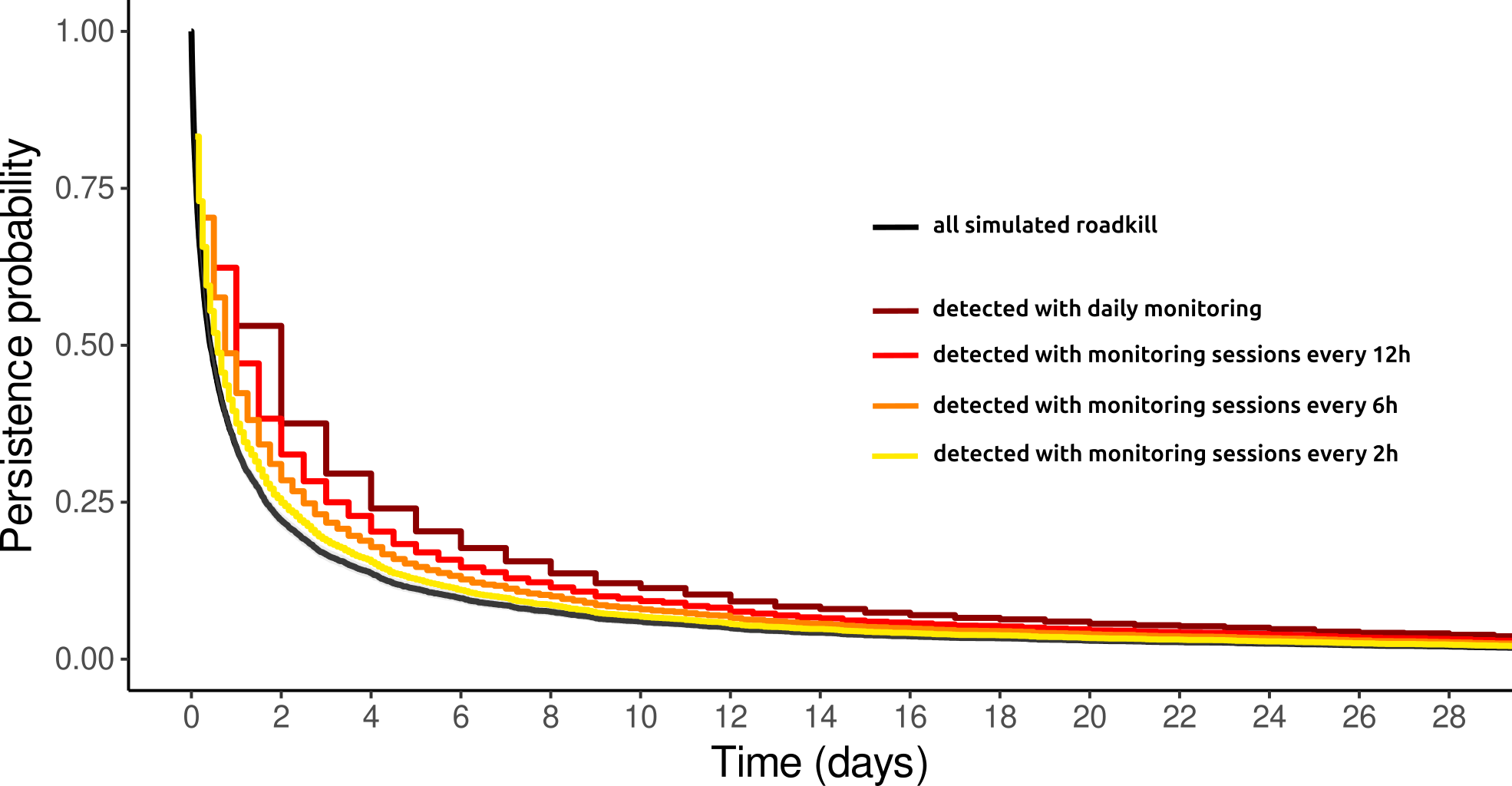
Figure A3: Kaplan-Meier survival curves for n=5000 simulated animal roadkill, for which the persistence on the road was randomly drawn from a log-normal distribution. The black curve represent the true persistence of roadkill for all simulated animals (median persistence: 38h18, 95% CI: 36h24,40h42). The remaining curves show the persistence estimated with experimental approaches where observers survey the road every 2, 6, 12 or 24h to detect new roadkill and subsequently record their persistence on the road. As the interval between surveys increases, the chances of missing carcasses with short persistence times increase, leading to an over-representation of long persistence times in the sampling process and thus, an overestimation of the persistence of roadkill. In this context, more regular surveys also allow for a finer resolution of the persistence times, leading to more precise estimates.
